# Bacteriocinogenic lactic acid bacteria in the traditional cereal-based beverage Boza: a genomic and functional approach

**DOI:** 10.1101/2021.10.05.463085

**Authors:** Luciano Lopes Queiroz, Christian Hoffmann, Gustavo Augusto Lacorte, Bernadette Dora Gombossy de Melo Franco, Svetoslav Dimitrov Todorov

## Abstract

Boza is a traditional low-alcohol fermented beverage from the Balkan Peninsula, frequently explored as a functional food product. The product is rich in Lactic Acid Bacteria (LAB) and some of them can produce bacteriocins. In this study, a sample of Boza from Belogratchik, Bulgaria, was analyzed for the presence of bacteriocinogenic LAB, and after analyses by RAPD-PCR, three representative isolates were characterized by genomic analyses, using whole genome sequencing. Isolates identified as *Pediococcus pentosaceus* ST75BZ and *Pediococcus pentosaceus* ST87BZ contained operons encoding for bacteriocins pediocin PA-1 and penocin A, while isolate identified as *P. acidilactici* ST31BZ contained only the operon for pediocin PA-1 and a CRISPR/Cas system for protection against bacteriophage infection. The antimicrobial activity of bacteriocins produced by the three isolates was inhibited by treatment of the cell-free supernatants with proteolytic enzymes. The produced bacteriocins inhibited the growth of *Listeria monocytogenes, Enterococcus* spp. and some *Lactobacillus* spp., among other tested species. The levels of bacteriocin production varied from 3200 AU/ml to 12800 AU/ml recorded against *L. monocytogenes* 104, 637 and 711, measured at 24 h of incubation at 37°C. All bacteriocins remained active after incubation at pH 2.0 to 10.0. The activity mode of the studied bacteriocins was bactericidal, as determined against *L. monocytogenes* 104, 637 and 711. In addition, bactericidal activity was demonstrated using a cell leakage β-galactosidase assay, indicating a pore formation mechanism as a mode of action. The present study highlights the importance of combining metagenomic analyses and traditional microbiological approaches as way of characterizing microbial interactions in fermented foods.

## Introduction

Boza is a traditional fermented cereal product, popular in many countries in the Balkan Peninsula, spread throughout the Middle East region by the Ottoman Empire^1,2^. There are several beneficial properties attributed to Boza: besides its high nutritional value, there is a diverse set of recommendations in traditional medicine^1^.

The microbiological composition of Boza includes several species of lactic acid bacteria (LAB), such as *Lactobacillus acidophilus, Lactobacillus brevis, Lactobacillus coprophilus, Lactobacillus coryniformis, Lactobacillus fermentum, Lactobacillus paracasei, Lactobacillus pentosus, Lactobacillus plantarum, Lactobacillus rhamnosus, Lactobacillus sanfrancisco, Lactococcus lactis* subsp. *lactis, Leuconostoc mesenteroides, Leuconostoc mesenteroides* subsp. *dextranicum, Leuconostoc raffinolactis, Leuconostoc lactis, Enterococcus faecium, Pediococcus pentosaceus, Oenococcus oeni, Weissella confusa* and *Weissella paramesenteriodes*^1,2^. Several of these LAB species contain beneficial strains, used as probiotics as starter cultures, and for food biopreservation^3^.

The application of LAB in food biopreservation is related to their ability of producing metabolites with antimicrobial activity, explored by traditional fermentation processes and traditional medicine for centuries. Bacteriocins constitute a group of inhibitory metabolites produced by LAB, presenting a polypeptide nature and produced in the bacterial ribosomal complex with no post-translational modification. Usually, the producer bacteria are immune to their own bacteriocins, due to mechanisms present in their genome^4^.

Bacteriocins have been described as effective against closely related species^4^. However, some reports have found unorthodox bacteriocin activities, such as inhibition of some viruses, *Mycobacterium spp*. as well as some fungi^5,6^. Several LAB isolated from Boza have been characterized for their ability to produce bacteriocins and proposed for different applications in biopreservation processes^1^.

Bacteriocins are known for their low cytotoxicity, being considered generally safe for human and animal consumption^7,8^. In the last decade, special attention was given to their pharmaceutical properties and potential application in human and veterinary medicine^4,5^.

In this study, we used a whole genome sequencing approach for the characterization of bacteriocins produced by some selected strains of lactic acid bacteria, isolated from Boza. This approach is important for a better understanding of the potential of application of this technological strategy to improve the safety of fermented food products.

## Results

### Screening for bacteriocinogenic LAB

The average bacterial population in the Boza samples was 1.1 × 10^8^ CFU/ml recorded by plating on MRS. Colony morphology on MRS agar plates, Gram-staining and catalase test indicated that the majority of the isolates were lactic acid bacteria (LAB), with coccoid and rod morphology. Preliminary screening tests for microbial inhibition indicated that most colonies were active against *Listeria monocytogenes* 104. The majority presented coccid morphology and were Gram positive, catalase negative and oxidase positive. The cell-free supernatants (CFS) of 18 isolates presented antimicrobial activity against *L. monocytogenes* 104 and *E. faecium* ATCC 10434. The activity was lost after treatment of the CFS with Proteinase K and pronase, but not when treated with α-amylase, lipase or catalase, indicating the proteinaceous nature of the antimicrobial activity. Bio-molecular fingerprinting based on RAPD-PCR grouped these isolates in 3 distinct groups. One isolate from each group was submitted to 16S rRNA sequencing for identification, resulting in *Pediococcus acidilactici* (one isolate) and *P. pentosaceus* (two isolates). These isolates were submitted to whole genome sequencing for a better characterization of the bacteriocin production.

### *Pediococcus acidilactici* ST31BZ genome characterization

The genome assembly for strain ST31BZ produced 19 contigs with N50 of 433,517 bp and 235x coverage. MiGA essential genes analysis showed a completeness of 94.6%, contamination of 1.8%, with an overall quality of 85.6%. We detected 105 of 111 essential genes in this genome and it was classified by MiGA using genome-aggregate Average Nucleotide and Amino Acid Identity (ANI/AAI) concepts as *P. acidilactici*, and closely related with *P. acidilactici* ZPA017 (NZ_CP015206.1) (98.89% ANI), isolated from black pig in Beijing, China.

The genome size for *P. acidilactici* ST31BZ isolate was 1,899,329 bp, with 42.18% GC-content, and 1,847 predicted protein-coding genes and no pseudogenes (Fig. 1a). We detected 50 tRNA and three rRNAs operons with complete 16S, 23S, and 5S coding regions. One CRISPR repeat with 36nt and 366 bp length was detected and a CAS-Type IIA System (*cas9, cas1, cas2*, and *csn2*) upstream of the CRISPR array. Two circular plasmids were found and designated as pST31BZ-1 with 49,093 bp and pST31BZ-2 with 9,090 bp containing a pediocin PA-1 operon (Fig. 2a). The pST31BZ-1 plasmid was similar to pHN9-1 (BLASTn, Identity: 98.19%; Coverage: 58%), a 42,239 bp plasmid found in *P. acidilactici* HN9 isolated from the Traditional Thai-Style Fermented Beef Nhang (Surachat *et al*., 2021). No antibiotic resistance genes were predicted using ARIBA in the genome or plasmids of strain ST31BZ, however, we were able to predict the following resistance genes using the KEGG GhostKoala automatic annotation pipeline: Lantibiotic tranporter system (NisE/F/G), Lincosamide and streptogramin A resistance (Lsa), Beta lactamase class A (PenP), Cationic antimicrobial peptide (CAMP) resistance, dltABCD operon (Dlt A/B/C/D), and two Multidrug efflux pump (EfrA/B and AbcA).

**Fig. 1.**
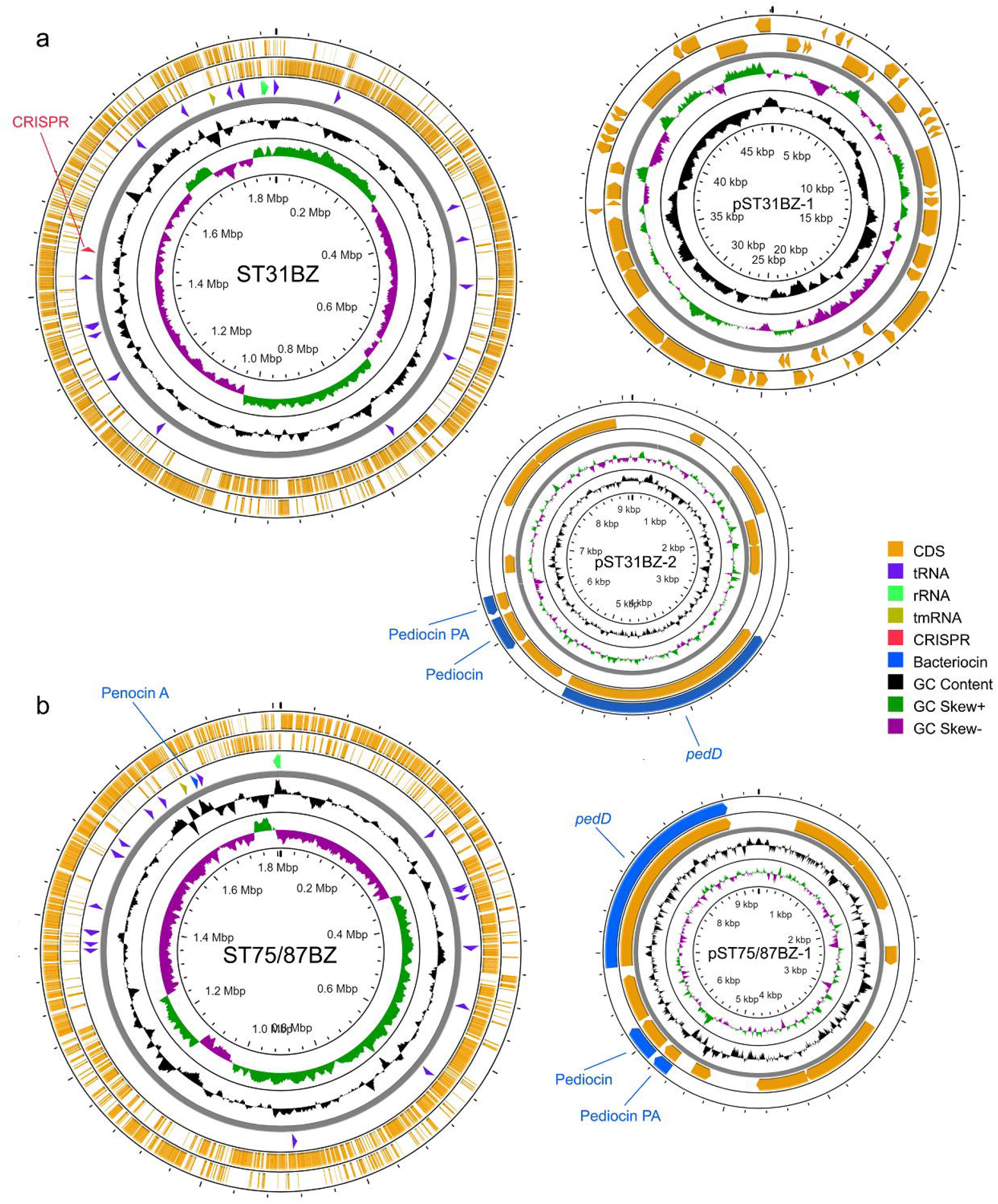
Genome and plasmids from two novel strains of LAB. **a**. *P. acidilactici* ST31BZ and its two plasmids. **b**. *P. pentosaceus* ST75BZ and ST87BZ and its plasmid. Representation is not drawn to scale to allow a good visual representation of the plasmid sequences.

**Fig. 2.**
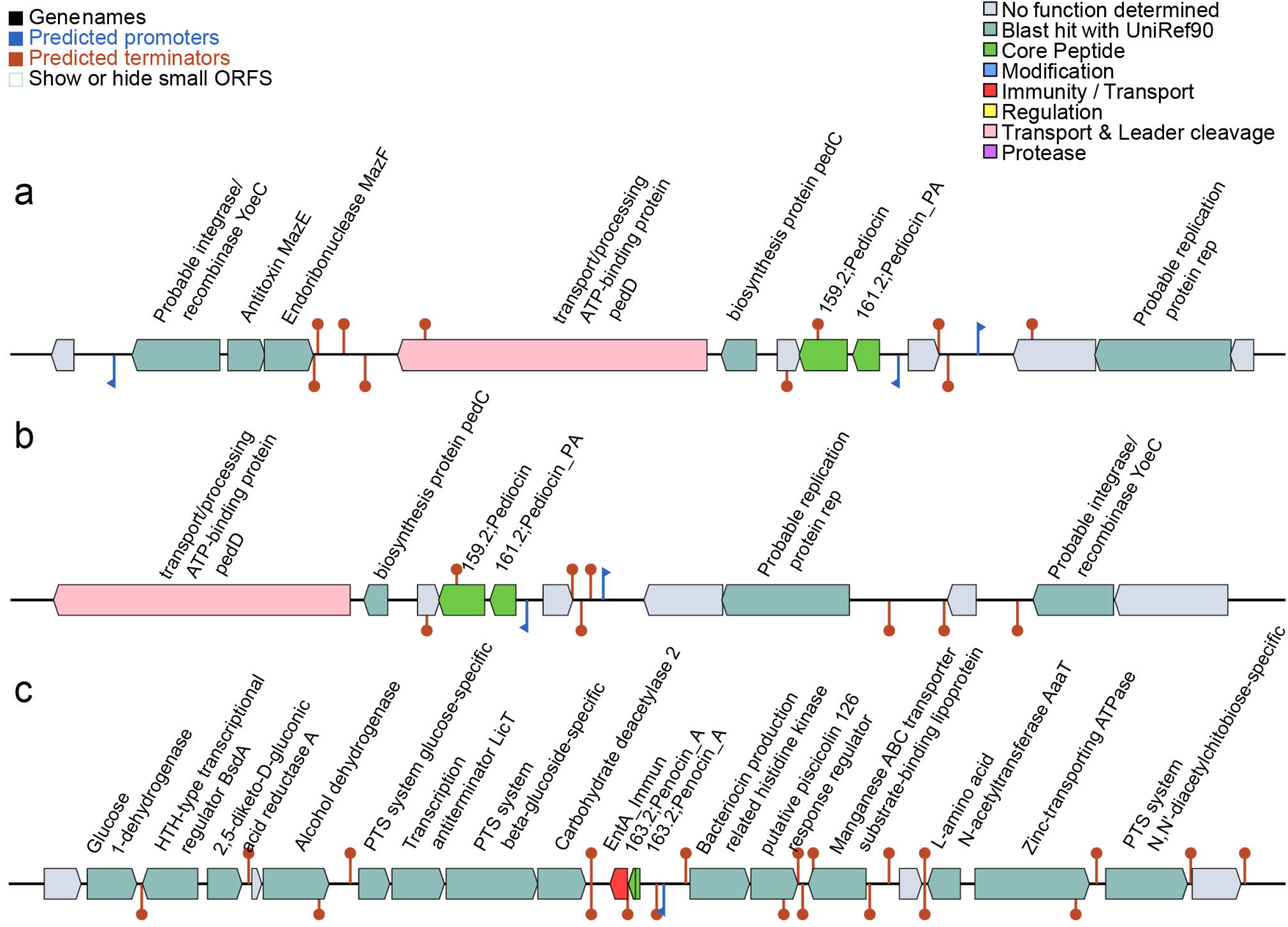
Putative bacteriocin gene clusters identified using BAGEL 4. **a**. Putative bacteriocin found pST31BZ-2 from *P. acidilactici* ST31BZ; **b**. Putative bacteriocin found pST75BZ and pST87BZ-1 from *Pediococcus pentosaceus* ST75BZ and ST87BZ; **c**. Putative bacteriocin found in genome of *Pediococcus pentosaceus* ST75BZ and ST87BZ.

*P. acidilactici* ST31BZ genome presented genes coding for several sugar transporters such as sucrose, maltose/glucose and cellobiose, and is likely able to degrade them all to lactate and acetate. We also found evidence that the strain is able to synthesize arginine, glycine, alanine, serine, asparagine, aspartate, glutamine and glutamate (Supplementary Table 1).

### *Pediococcus pentosaceus* ST75BZ and ST87BZ genome characterization

The assembled genomes of the two *P. pentosaceus* isolates presented a very high similarity (99.99% ANI), despite being analyzed separately, the only difference being that we obtained 15 contigs with N50 of 282,292 bp and 63x genome coverage for isolate ST75BZ, and 15 contigs with N50 of 296,046 bp and 62x genome coverage for isolate ST87BZ. The following results apply to both isolates and we will refer to them as strain *P. pentosaceus* ST75BZ and ST87BZ for the remainder of the text (Fig. 1b).

MiGA essential genes analysis showed a completeness of 95.5%, contamination of 1.8% and an excellent quality of 86.5% for both isolates. We detected 106 of 111 essential genes and this strain was taxonomically classified by MiGA, using genome-aggregate ANI/AAI concepts, as *P. pentosaceus*, and closely related with *P. pentosaceus* ATCC 25745 (NC_008525.1) (99.1% ANI). The genome size for *P. pentosaceus* ST75BZ and ST87BZ isolate was 1,810,333 bp, average GC content was 37.08%, with 1,773 protein-coding genes predicted and no pseudogenes. We detected 50 tRNA and three rRNAs operons with complete 16S, 23S, and 5S coding regions. No CRISPR repeats were detected. One circular plasmid was found and designated as pST75BZ and pST87BZ-1 with 9,342 bp containing a pediocin PA-1 operon (Fig. 2b). Other pediocin-like bacteriocin operon found was annotated as penocin A (Fig. 2c), present in the strain’s genome. We also found point mutations in the 23S rRNA coding sequence predicted to confer resistance to macrolides (azithromycin) and streptogramins using ARIBA. Using KEGG GhostKoala pipeline, other resistance genes were also detected: Beta lactamase class A gene (*PenP*), a Cationic Antimicrobial Peptide (CAMP) resistance gene, the dltABCD operon (Dlt A/B/C/D), two Multidrug efflux pump genes (*EfrA/B* and *AbcA*); and a lincosamide resistance gene (*Lsa*). The genome of *P. pentosaceus* ST75BZ and ST87BZ presented similar values for size, GC content, number of predicted proteins and ribosomal operons as 65 strains of *P. pentosaceus* in China^9^.

Strains *P. pentosaceus* ST75BZ and ST87BZ genome presented similar metabolic capabilities as *P. acidilacti* ST31BZ, however this strain has a reduced sugar transport capability, is able to synthesize lysine and is positive for genes encoding for a quorum sensing mechanism that we were not able to attribute to a distinct physiologic state or metabolic process (Supplementary Table 2).

### Pangenomic analysis of *Pediococcus* spp

Pangenomic analysis of the *Pediococcus* spp. isolates was carried out using Anvi’o v6.1 (Eren et al. 2015) with the pangenomic workflow. We selected five complete genomes of *P. acidilactici* and compared to *P. acidilactici* ST31BZ genome (Fig. 3a). *P. acidilactici* ST31BZ strain formed a related group with *P. acidilactici* ZPA017 (98.89% ANI) and BCC1 (98.84% ANI). We found 1487 core genes (1360 Single-copy Core genes, SCG) in all six genomes. *P. acidilactici* ST31BZ has 152 unique genes (Supplementary Table 3). COG annotation indicated prediction of six genes that encode phage-related proteins, in addition to six genes related to transposase activity, and 15 genes involved in carbohydrate metabolism, including a Na+/melibiose symporter and an arabinose efflux permease.

**Fig. 3.**
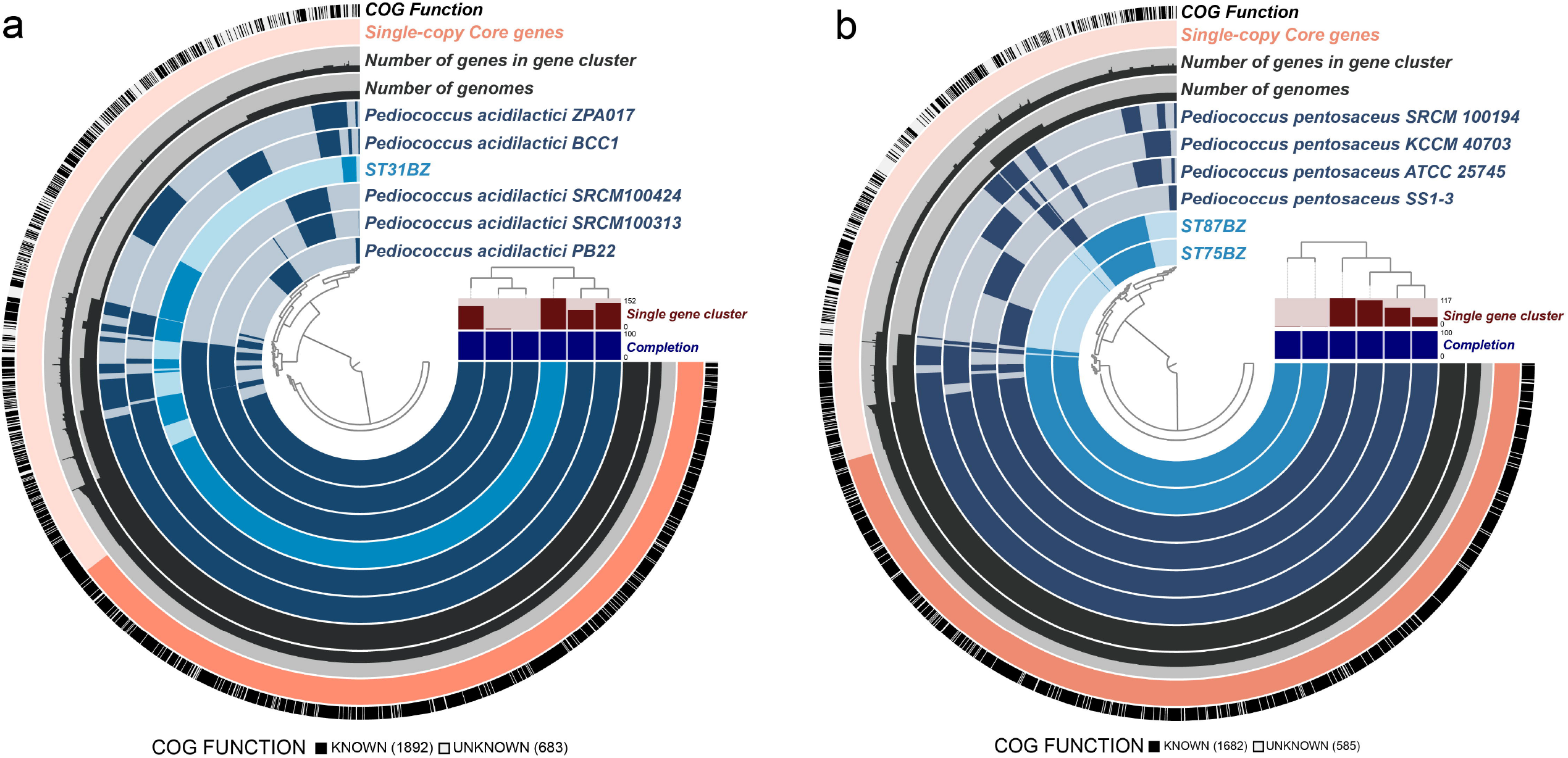
Pangenomic comparison. **a**. Isolate of *Pediococcus acidilactici* ST31BZ, **b**. *Pediococcus pentosaceus* ST75BZ and *Pediococcus pentosaceus* ST87BZ using Anvi’o. ST31BZ was compared with five complete genomes of *Pediococcus acidilactici* (strains ZPA017, BCC1, PB22, SRCM100424, and SRCM100313). ST75BZ and ST87BZ were compared with four complete genomes of *Pediococcus pentosaceus* (strains SS1-3, ATCC 25745, KCCM 40703, and SRCM 100194). Genomes of both pangenomes were organized based on gene cluster presence/absence.

Four complete genomes of *P. pentosaceus* were compared to the *P. pentosaceus* ST75/ST87 strain (Fig. 3b). *P. pentosaceus* ST75BZ and ST87BZ presented a total of 1446 core genes (1373 Single-copy Core genes, SCG) also observed in all compared genomes. However, *P. pentosaceus* ST75BZ and ST87BZ strains have 142 unique genes (Supplementary Table 4).

Based on the pangenomes of our isolates against reference genomes, we propose *P. pentosaceus* ST31BZ as a new strain of *P. acidilactici* and strains ST75BZ and ST87BZ as a new strain of *P. pentosaceus*. The two closest strains to *P. acidilactici* were ZPA017 and BCC-1, isolated from feces of a healthy pig^10^ and cecum of a broiler chicken^11^, respectively. The unique genes present in our isolates were related to antibiotic resistance and prophages, such as glycopeptide antibiotics resistance protein (COG4767) and phage portal protein BeeE (COG4695) in *P. acidilactici* ST31BZ (Supplementary Table 3); and ABC-type multidrug transport system (COG0842) and phage terminase large subunit (COG1783) in *P. pentosaceus* ST75BZ and ST87BZ. *P. pentosaceus* ST75BZ and ST87BZ, also showed the presence genes related to SOS response system, such as SOS-response transcriptional repressor (COG1974) (Supplementary Table 4).

Our two novel isolates presented pediocin PA-1 coding genes in their plasmids, which were highly similar (96% identity). Additionally, *P. pentosaceus* ST75BZ and ST87BZ presented a penocin A gene in its genome. Our annotation pipeline also detected bacteriocin genes in some of the strains used in the pangenome analysis: we detected two enterolyzin A genes in the genome of *P. acidilactici* BCC1, a bovicin coding gene in *P. acidilactici* BCC1 plasmid; an enterolysin A gene in the genome of *P. acidilactici* ZPA017; and a penocin A gene and enterolysin A gene in the genome of *P. pentosaceus* ATCC 25745.

### Bacteriocin production kinetics and stability tests

For *P. acidilactici* ST31BZ, *P. pentosaceus* ST75BZ and *P. pentosaceus* ST87BZ, only a small amount of bacteriocins were detected in the cell surfaces (approximately 200 AU/ml), with the majority of activity detected in cell-free culture supernatants. All remaining assays were carried out using cell-free supernatants, unless otherwise stated.

We characterized the bacteriocins produced by our three selected isolates for their stability in different pH, salt concentration, temperature and detergents. The three isolates grew well when cultured in MRS at 25, 30 and 37°C for 24 h, and produced bacteriocins (12800 AU/ml for ST31BZ and 3200 AU/ml for ST75BZ and ST87BZ). All further experiments were performed at 37°C, taking in consideration potential future applications of these strains as probiotics for human application. The activity of the bacteriocins remained unaltered after exposure to different pH (from 4.0 to 10.0), temperature (10, 25, 30, 37, 45, 80, 100 °C for up to 240 min, and at 121 °C for 15 min), or in the presence of NaCl, skim milk, SDS, Tween 20 and Tween 80 and EDTA, highlighting their potential versatility for use in industrial settings (data not shown).

The end-pH recorded for *P. acidilactici* ST31BZ, *P. pentosaceus* ST75BZ and *P. pentosaceus* ST87BZ when cultured overnight in MRS at 37°C were 4.43, 4.1 and 4.05, respectively (Fig. 4a). Over the same time period (27 h), the cell density increased from approximately OD_600nm_ 0.04 to 3.03 for *P. acidilactici* ST31BZ, 0.08 to 4.7 for *P. pentosaceus* ST75BZ and 0.06 to 4.77 for *P. pentosaceus* ST87BZ (Fig. 4a). Moreover, levels of bacteriocin produced by *P. acidilactici* ST31BZ were gradually increased during the fermentation process and reached 12800 AU/ml at 15 h and remained stable until the end of the monitored period of 27 h (Fig. 4a). However, bacteriocin produced by *P. pentosaceus* ST75BZ reached its maximum production levels (6400 AU/ml) at 15h from the beginning of fermentation, remained stable until 18h of fermentation time and decreased in the next 9 h (Fig. 4b). As expected, a similar bacteriocin production profile was recorded for both *P. pentosaceus* strains (Fig. 4c).

**Fig. 4.**
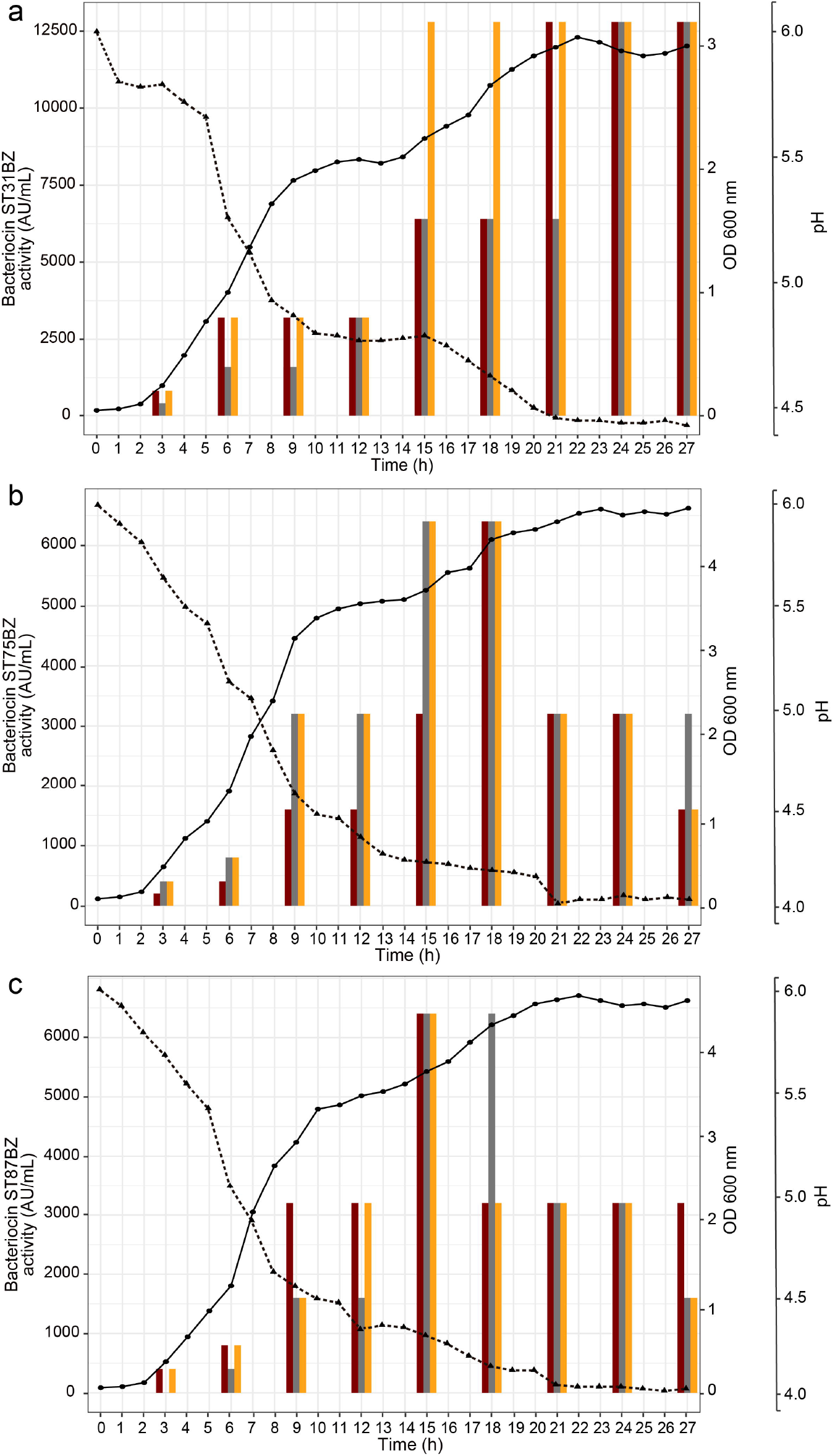
Bacterial growth. Growth changes were monitored as changes in OD_600 nm_ (-•-), pH (-▴-) and the production of bacteriocins evaluated (mean from 3 experiments is shown), against *Listeria monocytogenes* 104 (dark-red bar), *Listeria monocytogenes* 637 (grey bar) and *Listeria monocytogenes* 711 (mustard bar) expressed as AU/mL presented as histogram for **a**. *Pediococcus acidilactici* ST31BZ, **b**. *Pediococcus pentosaceus* ST75BZ and **c**. *Pediococcus pentosaceus* ST87BZ.

### Mode of action of bacteriocins produced by *P. pentosaceus*

The mechanisms of action of the bacteriocins produced by *P. acidilactici* ST31BZ, *P. pentosaceus* ST75BZ or *P. pentosaceus* ST87BZ was investigated using a growth inhibition assay, by adding 6400 AU/ml of bacteriocins produced by the strains to early-logarithmic growing (3-h-old) cultures of *L. monocytogenes* 104, 637 and 711. All three novel strains inhibited the growth of *Listeria* (Fig. 5), and recovery of viable *L. monocytogenes* from the cultures after 10 and 24 h of growth was not possible, indicating the killing effect.

**Fig. 5.**
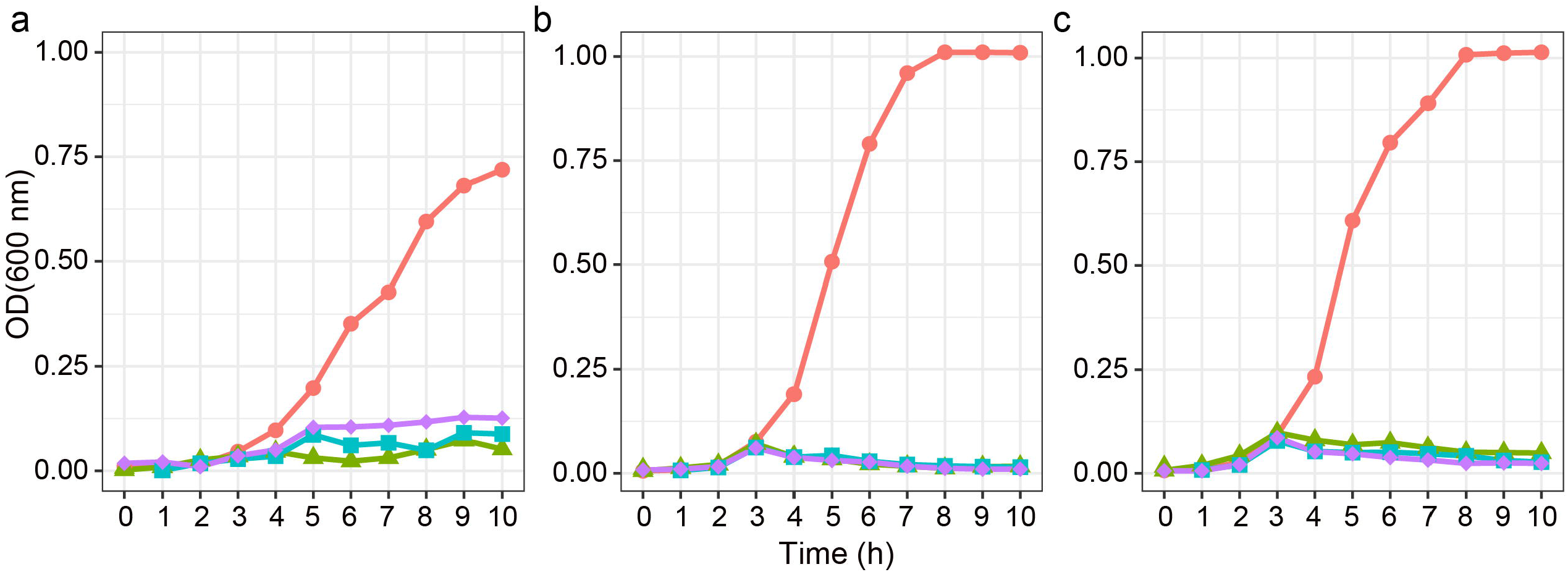
The effect of bacteriocin produced. Bacteriocin produced by *Pediococcus acidilactici* ST31BZ, *Pediococcus pentosaceus* ST75BZ and *Pediococcus pentosaceus* ST87BZ (mean from 3 experiments is shown), on the growth of **a**. *Listeria monocytogenes* 104, **b**. *Listeria monocytogenes* 637 and **c**. *Listeria monocytogenes* 711. Cultures received a standardized bacteirocin dose at 3h post culture start. Colors indicate the origin of the bacteriocin added: *P. acidilactici* ST31BZ - green (-▴-), *P. pentosaceus* ST75BZ blue (-◼-), and *P. pentosaceus* ST87BZ purple (-♦-). Negative control is shown in red (-•-).

We investigated cell membrane pore formation using a β-galactosidase leakage assay. Cells of *L. mono*cytogenes 104, 637 and 711 were incubated for 10 minutes with cell-free supernatants containing 6400 AU/ml of bacteriocin, followed by the addition of the β-galactosidase substrate and colorimetric product read-out. All bacteriocin/*Listeria* combinations tested produced a positive result, indicating that the mechanism of action involves pore formation and the destabilization of the cell membrane on the test strains.

### Bacteriocin activity spectrum

The cell-free culture supernatants of the three *Pediococcus* spp. was tested for activity against a panel of different species/strains. The growth of most *E. faecalis, E. faecium, E. hirae, Lc. lactis, L. innocua* and *L. monocytogenes*, some of *Lb. casei* and *Str. termophylus* strains was inhibited, indicating a similar spectrum of activity (Fig. 6, Supplementary Table 5).

**Fig. 6.**
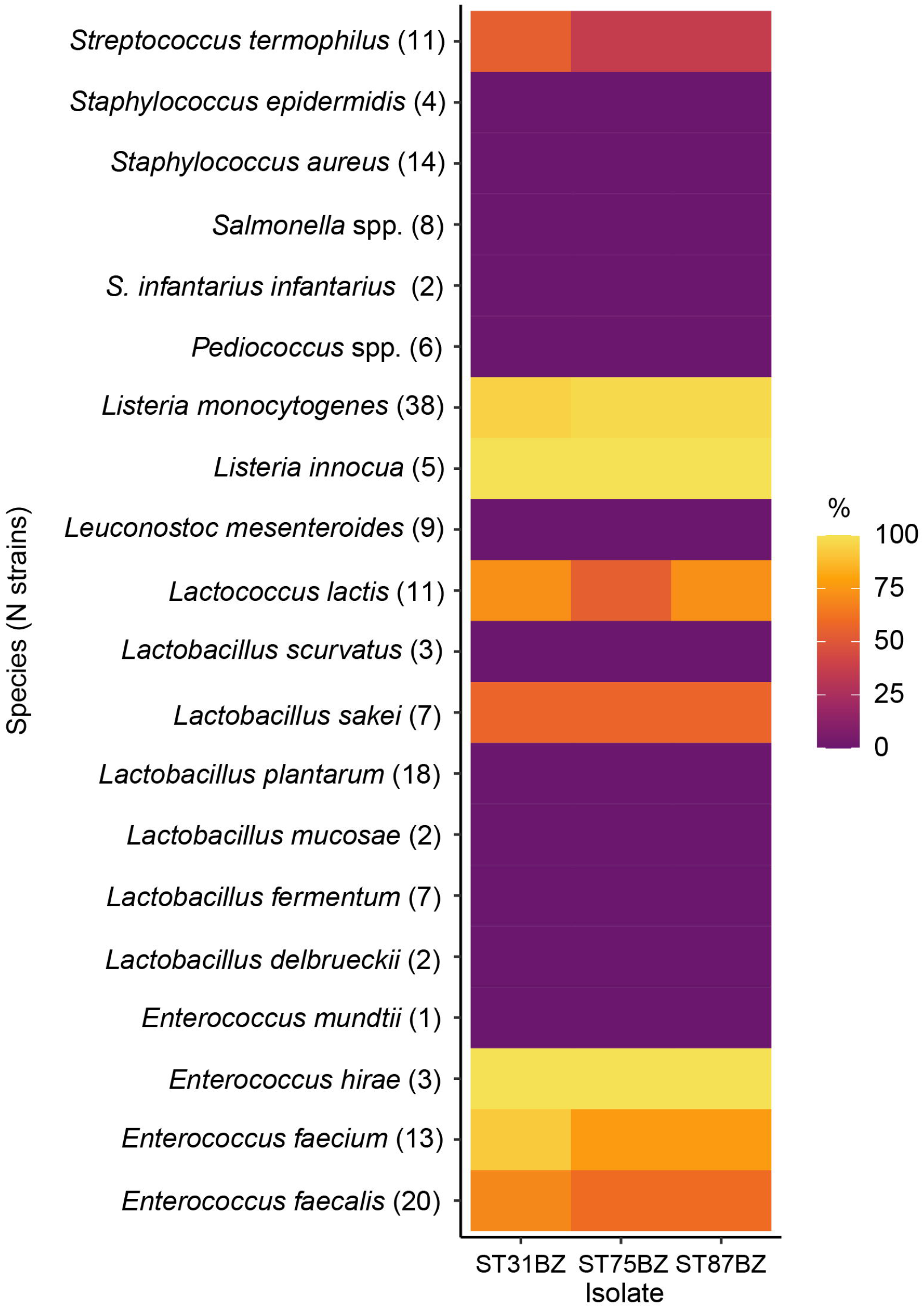
Bacterial susceptibility test results. The activity spectrum against tested strains is presented as a heatmap where colors indicate the proportion of susceptible strains from the total of tested strains (shown in parenthesis after the species name).

## Discussion

Boza is a traditional fermented beverage rich in nutrients, presenting a diverse microbial composition. Tests with Boza samples from different regions of the Balkan peninsula have shown that LAB and yeasts are the main microorganisms involved in the fermentation^1,2^. The present study aimed to characterize the potential the bacteriocin production by strains isolated from a sample of Boza obtained from a medium scale manufacture in North-west Bulgaria, using a mixed molecular and functional approach.

The screening method allowed us to select 3 bacterial isolates, initially classified as *Pediococcus* sp. Genome analysis indicated that these three isolates likely belong to two different *Pediococcus* species: one *P. acidilactii*, and two *P. pentosaceus*. The *P. acidilactici* ST31BZ genome showed similar values of size, GC content, number of predicted proteins and ribosomal operons as other *P. acidilactici* strains^10,12^. Detected CRISPR systems are similar to previously described in *P. acidilactici* HN9^13^. A small plasmid-encoded bacteriocin (∼ 9 kb) was detected, such as found in pSMB74^14^ and pCP289^15^. Several studies have shown that strains of *P. pentosaceus* are able to produce bacteriocins^5^, and in our study we observed the presence of two pediocin-like bacteriocins, pediocin PA-1 and penocin A, both of which have been previously reported in *P. pentosaceus* genomic analyses. Antibiotic resistance, prophages, plasmid and plasmid-associated bacteriocins have been indicated as source of variability at *P. pentosaceus* genomes^9^. Our *P. acidilactici* strain, however, also contains a plasmid-associated bacteriocin operon (pediocin PA-1), differing from other strains reported in the literature, such as ZPA017 and BCC-1^10,11^. The metabolic characteristics of the strains obtained indicated a wider metabolic potential for carbohydrate transport for the *P. acidilactii* transport while the *P. pentosaceus* presented he capability to synthesize lysine and had quorum sensing related genes.

Data from our initial screening indicated that the antimicrobial metabolites produced by the studied strains were proteinaceous in nature, which prompted us to further characterize the potential mechanisms of action and the activity range for the bacteriocins being produced. Although a peptide or protein molecule must be present for the antimicrobial activity to be detected, this observation does not preclude the possibility that other moieties may also be present in a larger complex that has the final bacteriocinogenic effect, as previously reported^16–18^. LAB can produce a variety of antimicrobial compounds, including diacetyl, hydrogen peroxide, carbon dioxide, low molecular antimicrobial substances, and organic acids such as phenyl lactic acid^16^. It is possible that other protein or peptide-derived antibiotics could be produced by these strains, as we have detected a gene signature for lantibiotic resistance in the genome of the *P. acidilactici* strain. However, based on the obtained results, we can clearly exclude the possibility of acid/s or H_2_O_2_ to be involved in the antimicrobial properties of the studied strains.

Identification of the genes involved in bacteriocin biosynthesis is an important part of the characterization of the antimicrobial agent as bacteriocin(s). Bacteriocin genes can be part of the bacterial genome, and their expression and detection in the cell free supernatant need to be confirmed. Therefore, a functional approach is important in evaluating the bacteriocin’s activity. Bacteriocins produced by the three selected strains presented a wide range of activity and were thermostable, maintaining activity after exposure to different temperatures up to 4 h, including at 121°C for 20 min. Such activity breadth is highly desirable for industrial applications. Most bacteriocins are molecules smaller than 10 kDa, although some can form complexes of higher molecular weight^19^. This small size confers stability in harsh conditions. Other reports have described that bacteriocins produced by different *Pediococccus* spp. are stable at different pH and after exposure to low and high temperatures^20^. Surfactants and salts normally have not been reported to negatively influence bacteriocins activity^21^, however, plantaricin C19 activity was affected by treatment with SDS or Triton X-100^22^. SDS did not cause loss of activity as has been observed for enterocin EJ97, bozacin B14 or bacteriocin ST194BZ^20,23^. However, pH can play a role on the stability of the bacteriocins, as been reported for leucocin F10^24^, Moreover, lactocin NK24 stability was decreased when exposed to 100 °C for 30 min and even was completely inactivated at 121 °C for 15 min^25^. Similar effect of temperature was reported for lactocin MMFII, produced by *L. lactis* MMFII^26^. Even nisin, one of the best studied bacteriocins, was shown to be inactivated after 15 min at 121 °C when incubated at pH 7.0, but not when incubated at pH 3.0^27^.

We observed a strong antimicrobial activity of *P. acidilactici* ST31BZ, *P. pentosaceus* ST75BZ and *P. pentosaceus* ST87BZ against a wide range of bacteria tested. These strains produce bacteriocins with strong antimicrobial activity recorded against all listeria strains and almost all enterococci evaluated. Nevertheless, no antimicrobial activity was observed against *S. aureus* and any *Lactobacillus* spp., *Leuconostoc* spp., *Samolella* spp. and other *Pediococcus* spp. strains tested. Specific anti-*Listeria* activity is an attribute reported for different studied pediocins and particularly pediocin PA-1. Even Heng *et al*.^19^ in his classification of bacteriocins, dedicated a special position for anti-*Listeria* pediocin-like bacteriocins. This specific activity was linked to the conservative amino-acid motive directly involved in the bacteriocin mode of action and pore formation process^19^.

There are reports in the literature that some bacteriocins can be adsorbed onto the producer’s cell surface, and this characteristic can be used to facilitate the purification process and/or increase the bacteriocin yield for industrial application^28^. In case of the studied bacteriocins, only low levels of cell-adsorption were recorded, which cannot be considered relevant in increasing production yield. Similar results have been shown for pediocins produced by *P. acidilactici* HA-6111-2 and HA-5692-3, respectively, against *L. innocua* N27 and *E. faecium* HKLHS^21^. Moreover, pediocin PA-1 has been reported as an effective bactericidal bacteriocin for the control of different *L. monocytogenes* strains, leading to is proposition as prospective biopreservation for different fermented food products^21,29,30^. Furthermore, the results obtained from the β-galactosidase leakage assay confirm that bacteriocins produced by *P. acidilactici* ST31BZ, *P. pentosaceus* ST75BZ and *P. pentosaceus* ST87BZ likely destabilize the cell membrane inducing the cytoplasmic leakage and a complete loss of viability for the susceptible strains (data not shown). Similar approach and results have been reported for bacteriocins produced by *Lb. plantarum, Le. lactis* and *E. faecium*^16^, *Lb. buchneri*^31^, *Lb. plantarum*^32,33^ and *Lc. lactis* subsp. *lactis*^34^.

## Conclusion

This study confirm previous ones that have shown boza to be a rich source of LAB of biotechnological and industrial interest. The bacteriocinogenic potential of the isolated *P. acidilactici* ST31BZ, *P. pentosaceus* ST75BZ and *P. pentosaceus* ST87BZ strains reported here include a partial characterization of the mechanism of action and breadth of susceptible bacterial targets. As most of the fermented traditional food products, Boza is a rich multimicrobial system, were interactions between different microbial species is essential for the final product’s characteristics. Results of whole genome sequencing analysis of the three selected bacteriocinogenic strains, coupled with functional assays, highlight their metabolic potential for industrial application, including production at industrial level of Boza or any other fermented functional food product.

## Materials and methods

### Screening for bacteriocinogenic lactic acid bacteria in boza

Boza samples were obtained from a medium-scale manufacturer in North-west Bulgaria. Isolation of bacteriocinogenic lactic acid bacteria (LAB) from these samples was done according to the 3-layer method^35^. Samples of boza were submitted to serial decimal dilutions in sterile saline (0.85% NaCl, Sigma Diagnostics, St. Louis, MO, USA) and plated on the surface of MRS agar (Difco BD, Franklin Lakes, NJ, USA) plates. After addition of a layer of 1.0% agar (Difco), the plates were incubated for 48 h at 37°C. The colonies were counted and plates with well isolated colonies were added of an extra layer of BHI agar (Difco) (BHI supplemented with 1.0% agar) containing *L. monocytogenes* 104 or *E. faecium* ATCC 19434 (10^5^ CFU/ml). Plates were incubated for additional 24 h and colonies with visible inhibition zones were transferred to new BHI and incubated at 37 °C for 24 h. The cultures were checked for purity by streaking on MRS agar. Individual colonies on MRS agar were submitted to Gram-staining and catalase and oxidase tests. Presumed bacteriocin producing LAB and other microorganisms used as target organisms were stored at -80 °C in MRS or BHI added of 20% glycerol.

### Identification of the antimicrobial metabolite produced by the selected strains

The isolates were grown in MRS broth at 37 °C for 24 h, and submitted to centrifugation (6 000 xg, 10 min, 4 °C) for obtention of cell free supernatants (CFS). After adjustment of the pH to 5.0-6.5 with 1M NaOH, the CFS were heated for 10 min at 80°C to eliminate potential inhibitory effect of organic acids and to inactivate hydrogen peroxide and proteolytic enzymes in the medium. Ten microliters of treated CFS were spotted on the surface on plates containing BHI supplemented with 1.0% agar plates containing 10^5^ CFU/ml *L. monocytogenes* 104 or *E. faecium* ATCC 19434, used as target test micro-organisms. Plates were incubated at 37°C for 24 h and presence of growth inhibition zones wider than 2 mm was considered evidence for potential bacteriocin production (Supplementary Table 6)

The proteinaceous nature of the antimicrobial substances produced by the isolates was investigated in the cell free supernatants by treatment with Proteinase K, pepsin and pronase (all from Sigma), as described before^35^. Antimicrobial activity and the effect of temperature (30, 60, 120 and 240 min at 8, 25, 30, 37, 45, 60, 80 and 100°C, and 20 min at 121°C), pH (2.0, 4.0, 6.0, 8.0 and 10.0) and selected chemicals (NaCl, SDS, Tween 20, and Tween 80) on stability of produced bacteriocins were evaluated according dos Santos *et al*.^35^. Tests were performed with *L. monocytogenes* 104 and *E. faecium* ATCC 19434 as targets. All experiments were performed at least in duplicate in two independent occasions.

### Identification of isolates

The strains were identified based on recommended morphological, biochemical and physiological tests, according the Bergey’s Manual^36^ and 16S rRNA partial gene sequencing. Cultures were prepared in 50 ml MRS broth for 24 h at 37°C, cells collected by centrifugation (6000 x*g*, 10 min, 4°C) and DNA extracted using the ZR Fungal/Bacterial DNA Kit (Zymo Research, Irvine, CA, USA), following the manufacturer’s protocol. The obtained DNA was quantified using a NanoDrop (Thermo Fisher Scientific, Waltham, MA, USA). RAPD-PCR with Primers OPL-14 (5’-GTG ACA GGC T-3’) and OPL-20 (5’-TGG TGG ACC A-3’) was used for differentiation between the studied isolates^37^. Based on the RAPD profiles, three strains were selected and their DNAs subjected to PCR to amplify a region of 16S rRNA^37^, and further sequenced in the Center for Human Genome Studies (Institute of Biomedical Sciences, University of São Paulo, São Paulo, SP, Brazil). For identifications, the obtained sequences were analyzed in the Basic Local Alignment Search Tool (BLAST, GenBank, National Center for Biotechnology Information, Bethesda, MD, USA).

### Genomic analysis

Three isolates were selected for Whole Genome Sequencing. The DNA was extracted from cultures grown in MRS broth at 37°C for 24 h, using the ZR Fungal/Bacterial DNA Kit (Zymo Research). Extracted DNA was used for library preparation with Nextera DNA Flex Library Prep Kit (Illumina, San Diego, CA, USA). Libraries were sequenced on NextSeq Genome Sequencer (Illumina) with a NextSeq 500/550 Mid Output Kit v2.5 at Core Facility for Scientific Research – University of Sao Paulo (USP) (CEFAP-USP).

The obtained reads were processed with the software package BBTools (https://jgi.doe.gov/data-and-tools/bbtools/). Reads were trimmed for Nextera adapters and filtered to have an average Q-score < 15. Trimmed reads from the three strains were assembled *de novo* with SPAdes 3.13.0^38^. ORFs were predicted for all assembled contigs with Prokka pipeline^39^. Scaffolds were uploaded to MiGA to access genome assembly completeness and to identify the closest bacterial strain to each assembled genome^40^. tRNA were detected with ARAGORN^41^ and rRNAs with Barrnap (https://github.com/tseemann/barrnap). CRISPR regions were detected with PILER-CR^42^ and CRISPRCasFinder^43^. Plasmid searches were made using metagenomic plasmid function (metaplasmidspades.py)^44^ and the presence of plasmid genes verified using the script viralVerify^45^ with -p argument. Bacteriocins genes were detected with BAGEL4^46^ and antibiotic resistances genes using ARIBA^47^ with quality filtered reads and CARD database^48^.

Pangenomic analysis of *Pediococcus* isolates were done using anvi’o v6.1^49^ with the pangenomic workflow. Our objective was to compare the similarity of isolates and reference genomes. We focused on *P. acidilactici* and *P. pentosaceus* species. All sequencing data generated in this study can be accessed from GenBank Database at BioProject PRJNA731169. The genome of isolate ST31BZ was compared to five reference genomes of *P. acidilactici* species available on NCBI (BioProject numbers: PRJNA312971, PRJNA357663, PRJNA422477, PRJNA386762, and PRJNA386761). The genomes of isolates ST75BZ and ST87BZ were compared to four references genomes *of P. pentosaceus* species available on NCBI (PRJNA399825, PRJNA398, PRJNA376813, and PRJNA390207).

### Spectrum of activity

The selected bacteriocinogenic strains were grown in MRS at 37°C for 24 h and cell free supernatants were obtained as described before. Inhibitory effectiveness of the produced bacteriocins was evaluated against several LAB, selected food borne pathogens and some Gram-negative organisms, listed in Supplementary Table 5. The growth conditions (culture medium and temperature) were according to the recommended for each test microorganism. Test microorganisms were grown overnight and incorporated in appropriate medium, supplemented with 1.0% agar, at final concentration around 10^5^ CFU/ml. Studied bacteriocins were spotted (10 μl) on the surface and plates cultured for 24 h at recommended growth temperature. Zones of inhibition, larger than 2 mm were considered as positive result.

### Growth and bacteriocin production dynamics

The selected strains were grown in 20 ml MRS broth (Difco) at 25 °C, 30 °C or 37 °C for 24 h. Cell free supernatant was prepared as described before and bacteriocin activity determined against *L. monocytogenes* 104 and expressed in AU/ml. After selection the optimal temperature for bacteriocin production, the dynamic of the production was evaluated as follows: overnight cultures were prepared in 300 mL MRS broth (Difco) at 37°C, and optical density at 600 nm and pH changes were monitored hourly for 24 h. Production of bacteriocin(s) was measuring the antimicrobial activity against *L. monocytogenes* 104, 637 and 711 every three hours (expressed as AU/ml). Experiments were performed in two independent occasions.

### Inhibitory effect of bacteriocins evaluated via cell lysis assay

For evaluation of the effect of studied bacteriocin on actively growing *L. monocytogenes* 104, 637 and 711, the approach proposed by de Castilho *et al*.^50^ was followed. For the experiment, 300 ml BHI broth was inoculated with 1% (v/v) of *L. monocytogenes* 104, 637 or 711 and incubated for 3 h at 37°C. 30 ml filter-sterilized (0.22 μm Millipore sterile filters, Burlington, MA, USA) cell-free supernatant of each bacteriocinogenic strain was added to the culture and changes on the turbidity were monitored at OD_600nm_ every hour for 10 h. In addition, bacterial growth was determined after 24 h of incubation, to check for viable cells. The numbers of CFU/ml were determined after 10 h and 24 h of cultivation for all assays, i.e., with added bacteriocin and controls (without added bacteriocin), by plating on BHI supplemented with 2% agar and incubation at 37°C for 48 h. Experiments were performed in duplicate in 2 independent occasions.

### β-galactosidase assay

The capability of bacteriocin(s) to induce pore formation in the target strains was tested determining the level of β-galactosidase secreted from damaged cells. Cells from 20 ml of actively growing, log-phase cultures of *L. monocytogenes* 104, 637 and 711 were harvested, washed twice with 20 mL 0.03 M sodium phosphate buffer (K_2_HPO_4_/KH_2_PO_4_, pH 6.5) and the pellet re-suspended into 10 ml of the same buffer. Two ml of the cell suspensions were treated with equal volumes of each studied bacteriocin to yield final concentrations of 6 400 AU/ml. After 10 min at 37°C, 0.2 mL 0.1 M ONPG (O-nitrophenyl-β-D-galactopyranoside, Sigma), dissolved in 0.03 M sodium phosphate buffer (pH 6.8), was added to each of the cell suspensions and the cells incubated for additional 10 min at 37°C. The β-galactosidase reaction was stopped by adding 2.0 ml 0.1 M sodium carbonate. The cells were harvested by centrifugation (10 000 x *g*, 15 min, 25°C) and absorbance readings of cell-free supernatants recorded at 420 nm. Cultures of *L. monocytogenes* 104, 637 and 711 not treated with the studied bacteriocins were used as controls.

### Adsorption to cell surface of producer

The capability of the studied bacteriocins to adsorb to the producer cells was tested according to Yang *et al*.^28^, using *L. monocytogenes* 104. Experiments were performed in a triplicate.

### Vizualization

Circular representations of the isolate genomes were made using CGView Server^51^. The figures were made using the statistical software R^52^, with package ggplot2^53^.

## Supporting information

Supplemental informations

## Data Availability

Genomes assembled and their SRA sequencing data are available in the NCBI under BioProject PRJNA731169.

## Acknowledgements

We would like to thank all members of the Food Microbiology Laboratory for their support during the experiments. This work was supported by the Sao Paulo Research Foundation (FAPESP), grant: 2013/07914-8, and the Coordenação de Aperfeiçoamento de Pessoal de Nível Superior - Brasil (CAPES) - Finance Code 001.

## Contributions

Microbiology was performed by S.D.T. Sample processing and sequencing were performed by L.L.Q and G.A.L. Genome and statistical analyses were performed by L.L.Q and C.H. Manuscript was prepared by L.L.Q, G.A.L. C.H., and S.D.T. Study design was performed by C.H., and S.D.T. Contributed funding was made by B.D.G.M.F., C.H., and S.D.T. All authors reviewed and approved the manuscript.

## References

1. Leblanc, J. G. & Todorov, S. D. Bacteriocins producing lactic acid bacteria isolated from Boza, a traditional fermented beverage from Balkan Peninsula – from isolation to application. in Science against microbial pathogens: communicating current research and technological advances (ed. Méndez-Vilas, A.) 1311–1320 (Formatex Research Center, 2011).

2. Todorov, S. D. & Holzapfel, W. H. Traditional cereal fermented foods as sources of functional microorganisms. in Advances in Fermented Foods and Beverages 123–153 (Elsevier, 2015). doi:10.1016/B978-1-78242-015-6.00006-2

3. Marco, M. L. et al. The International Scientific Association for Probiotics and Prebiotics (ISAPP) consensus statement on fermented foods. Nat. Rev. Gastroenterol. Hepatol. 18, 196–208 (2021).

4. Chikindas, M. L., Weeks, R., Drider, D., Chistyakov, V. A. & Dicks, L. M. Functions and emerging applications of bacteriocins. Curr. Opin. Biotechnol. 49, 23–28 (2018).

5. Todorov, S. D., de Melo Franco, B. D. G. & Tagg, J. R. Bacteriocins of Gram-positive bacteria having activity spectra extending beyond closely-related species. Benef. Microbes 10, 315–328 (2019).

6. Todorov, S. D. & Chikindas, M. L. Lactic Acid Bacteria Bacteriocins and their Impact on Human Health. in Lactic Acid Bacteria (eds. Albuquerque, M. A. C., LeBlanc, A., LeBlanc, J. & Bedani, R.) 79–92 (CRC Press,2020). doi:10.1201/9780429422591-5

7. Wachsman, M. B. et al. Enterocin CRL35 inhibits late stages of HSV-1 and HSV-2 replication in vitro. Antiviral Res. 58, 17–24 (2003).

8. Torres, N. I. et al. Safety, Formulation and In Vitro Antiviral Activity of the Antimicrobial Peptide Subtilosin Against Herpes Simplex Virus Type 1. Probiotics Antimicrob. Proteins 5, 26–35 (2013).

9. Jiang, J. et al. Comparative Genomics of Pediococcus pentosaceus Isolated From Different Niches Reveals Genetic Diversity in Carbohydrate Metabolism and Immune System. Front. Microbiol. 11, 1–22 (2020).

10. Liu, H. et al. Effects of dietary supplementation with Pediococcus acidilactici ZPA017 on reproductive performance, fecal microbial flora and serum indices in sows during late gestation and lactation. Asian-Australasian J. Anim. Sci. 33, 120–126 (2020).

11. Lei, Z. et al. Transcriptomic Analysis of Xylan Oligosaccharide Utilization Systems in Pediococcus acidilactici Strain BCC-1. J. Agric. Food Chem. 66, 4725–4733 (2018).

12. Porto, M. C. W., Kuniyoshi, T. M., Azevedo, P. O. S., Vitolo, M. & Oliveira, R. P. S. Pediococcus spp.: An important genus of lactic acid bacteria and pediocin producers. Biotechnol. Adv. 35, 361–374 (2017).

13. Surachat, K., Kantachote, D., Deachamag, P. & Wonglapsuwan, M. Genomic Insight into Pediococcus acidilactici HN9, a Potential Probiotic Strain Isolated from the Traditional Thai-Style Fermented Beef Nhang. Microorganisms 9, (2021).

14. Motlagh, A., Bukhtiyarova, M. & Ray, B. Complete nucleotide sequence of pSMB 74, a plasmid encoding the production of pediocin AcH in Pediococcus acidilactici. Lett. Appl. Microbiol. 18, 305–312 (1994).

15. Kaur, B. & Balgir, P. Pediocin CP2GENE Localisation To Plasmid PCP289 Of <italic>Pediococcus Acidilactici</italic> MTCC 5101. Internet J. Microbiol. 3, 1–7 (2012).

16. Todorov, S. D. Diversity of bacteriocinogenic lactic acid bacteria isolated from boza, a cereal-based fermented beverage from Bulgaria. Food Control 21, 1011– 1021 (2010).

17. Lewus, C. B., Sun, S. & Montville, T. J. Production of an Amylase-Sensitive Bacteriocin by an Atypical Leuconostoc paramesenteroides Strain. Appl. Environ. Microbiol. 58, 143 (1992).

18. Keppler, P., Holz, U., Thielemann, F. W. & Meinig, R. Locked Posterior Dislocation of the Shoulder: Treatment Using Rotational Osteotomy of the Humerus. J. Orthop. Trauma 8, 286–292 (1994).

19. Heng, N. C. K., Wescombe, P. A., Burton, J. P., Jack, R. W. & Tagg, J. R. The Diversity of Bacteriocins in Gram-Positive Bacteria. in Bacteriocins 45–92 (Springer Berlin Heidelberg,2007). doi:10.1007/978-3-540-36604-1_4

20. Todorov, S. D. & Dicks, L. M. T. Pediocin ST18, an anti-listerial bacteriocin produced by Pediococcus pentosaceus ST18 isolated from boza, a traditional cereal beverage from Bulgaria. Process Biochem. 40, 365–370 (2005).

21. Albano, H. et al. Characterization of two bacteriocins produced by Pediococcus acidilactici isolated from “Alheira”, a fermented sausage traditionally produced in Portugal. Int. J. Food Microbiol. 116, 239–247 (2007).

22. Atrih, A., Rekhif, N., Moir, A. J. ., Lebrihi, A. & Lefebvre, G. Mode of action, purification and amino acid sequence of plantaricin C19, an anti-Listeria bacteriocin produced by Lactobacillus plantarum C19. Int. J. Food Microbiol. 68, 93–104 (2001).

23. Gálvez, A. et al. Isolation and characterization of enterocin EJ97, a bacteriocin produced by Enterococcus faecalis EJ97. Arch. Microbiol. 171, 59–65 (1998).

24. Parente, E., Moles, M. & Ricciardi, A. Leucocin F10, a bacteriocin from Leuconostoc carnosum. Int. J. Food Microbiol. 33, 231–243 (1996).

25. Lee, N.-K. & Paik, H.-D. Partial characterization of lacticin NK24, a newly identified bacteriocin of Lactococcus lactis NK24 isolated from Jeot-gal. Food Microbiol. 18, 17–24 (2001).

26. Ferchichi, M., Frère, J., Mabrouk, K. & Manai, M. Lactococcin MMFII, a novel class IIa bacteriocin produced by Lactococcus lactis MMFII, isolated from a Tunisian dairy product. FEMS Microbiol. Lett. 205, 49–55 (2001).

27. Noonpakdee, W., Santivarangkna, C., Jumriangrit, P., Sonomoto, K. & Panyim, S. Isolation of nisin-producing Lactococcus lactis WNC 20 strain from nham, a traditional Thai fermented sausage. Int. J. Food Microbiol. 81, 137–45 (2003).

28. Yang, R., Johnson, M. C. & Ray, B. Novel method to extract large amounts of bacteriocins from lactic acid bacteria. Appl. Environ. Microbiol. 58, 3355 (1992).

29. Loessner, M., Guenther, S., Steffan, S. & Scherer, S. A Pediocin-Producing Lactobacillus plantarum Strain Inhibits Listeria monocytogenes in a Multispecies Cheese Surface Microbial Ripening Consortium. Appl. Environ. Microbiol. 69, 1854–1857 (2003).

30. Osmanagaoglu, O., Kiran, F. & Nes, I. F. A probiotic bacterium, Pediococcus pentosaceus OZF, isolated from human breast milk produces pediocin AcH/PA-1. African J. Biotechnol. 10, 2070–2079 (2011).

31. Yıldırım, Z., Avşar, Y. K. & Yıldırım, M. Factors affecting the adsorption of buchnericin LB, a bacteriocin produced by Lactocobacillus buchneri. Microbiol. Res. 157, 103–107 (2002).

32. Todorov, S. D. & Dicks, L. M. T. Screening for bacteriocin-producing lactic acid bacteria from boza, a traditional cereal beverage from Bulgaria: Comparison of the bacteriocins. Process Biochem. 41, 11–19 (2006).

33. Bhunia, A. K., Johnson, M. C., Ray, B. & Kalchayanand, N. Mode of action of pediocin AcH from Pediococcus acidilactici H on sensitive bacterial strains. J. Appl. Bacteriol. 70, 25–33 (1991).

34. Todorov, S. D., Botes, M., Danova, S. T. & Dicks, L. M. T. Probiotic properties of Lactococcus lactis ssp. lactis HV219, isolated from human vaginal secretions. J. Appl. Microbiol. 103, 629–639 (2007).

35. dos Santos, K. M. O. et al. Exploring Beneficial/Virulence Properties of Two Dairy-Related Strains of Streptococcus infantarius subsp. infantarius. Probiotics Antimicrob. Proteins 12, 1524–1541 (2020).

36. De Vos, P. et al. Bergey’s Manual of Systematic Bacteriology. (Wiley Publishing Group, 2009).

37. de Moraes, G. M. D. et al. Functional Properties of Lactobacillus mucosae Strains Isolated from Brazilian Goat Milk. Probiotics Antimicrob. Proteins 9, 235–245 (2017).

38. Bankevich, A. et al. SPAdes: A New Genome Assembly Algorithm and Its Applications to Single-Cell Sequencing. J. Comput. Biol. 19, 455–477 (2012).

39. Seemann, T. Prokka: Rapid prokaryotic genome annotation. Bioinformatics 30, 2068–2069 (2014).

40. Rodriguez-R, L. M. et al. The Microbial Genomes Atlas (MiGA) webserver: Taxonomic and gene diversity analysis of Archaea and Bacteria at the whole genome level. Nucleic Acids Res. 46, W282–W288 (2018).

41. Laslett, D. & Canback, B. ARAGORN, a program to detect tRNA genes and tmRNA genes in nucleotide sequences. Nucleic Acids Res. 32, 11–16 (2004).

42. Edgar, R. C. PILER-CR: Fast and accurate identification of CRISPR repeats. BMC Bioinformatics 8, 1–6 (2007).

43. Couvin, D. et al. CRISPRCasFinder, an update of CRISRFinder, includes a portable version, enhanced performance and integrates search for Cas proteins. Nucleic Acids Res. 46, W246–W251 (2018).

44. Antipov, D., Raiko, M., Lapidus, A. & Pevzner, P. A. Plasmid detection and assembly in genomic and metagenomic data sets. Genome Res. 29, 961–968 (2019).

45. Antipov, D., Raiko, M., Lapidus, A. & Pevzner, P. A. Metaviral SPAdes: Assembly of viruses from metagenomic data. Bioinformatics 36, 4126–4129 (2020).

46. van Heel, A. J. et al. BAGEL4: a user-friendly web server to thoroughly mine RiPPs and bacteriocins. Nucleic Acids Res. 46, W278–W281 (2018).

47. Hunt, M. et al. ARIBA: Rapid antimicrobial resistance genotyping directly from sequencing reads. Microb. Genomics 3, 1–11 (2017).

48. Alcock, B. P. et al. CARD 2020: Antibiotic resistome surveillance with the comprehensive antibiotic resistance database. Nucleic Acids Res. 48, D517–D525 (2020).

49. Eren, A. M. et al. Anvi’o: an advanced analysis and visualization platform for ‘omics data. PeerJ 3, e1319 (2015).

50. Castilho, N. P. A., Colombo, M., Oliveira, L. L. de, Todorov, S. D. & Nero, L. A. Lactobacillus curvatus UFV-NPAC1 and other lactic acid bacteria isolated from calabresa, a fermented meat product, present high bacteriocinogenic activity against Listeria monocytogenes. BMC Microbiol. 19, 63 (2019).

51. Stothard, P., Grant, J. R. & Van Domselaar, G. Visualizing and comparing circular genomes using the CGView family of tools. Brief. Bioinform. 20, 1576– 1582 (2018).

52. R Development Core Team. R: A language and environment for statistical computing. (2020).

53. Wickham, H. ggplot2. (Springer International Publishing, 2016). doi:10.1007/978-3-319-24277-4

