## Supplemental informations for "Bacteriocinogenic lactic acid bacteria in the traditional cereal-based beverage Boza: a genomic and functional approach"

Supplementary Table 5. Bacteriocin susceptibility assay results, including species name and number of strains tested.

| Test organisms | growth media | Bacteriocin activity  sensitive strains / total number tested strains | | | | | |
| --- | --- | --- | --- | --- | --- | --- | --- |
|  |  | ST31BZ | | ST75BZ | | | ST87BZ |
| *Enterococcus faecalis* | MRS | 14 / 20 | 12 / 20 | | | 12 / 20 | |
| *Enterococcus faecium* | MRS | 12 / 13 | 10 / 13 | | | 10 / 13 | |
| *Enterococcus hirae* | MRS | 3 / 3 | 3 / 3 | | | 3 / 3 | |
| *Lactococcus lactis* | MRS | 8 / 11 | 6 / 11 | | | 8 / 11 | |
| *Listeria innocua* | BHI | 5 / 5 | 5 / 5 | | | 5 / 5 | |
| *Listeria monocytogenes* | BHI | 36 / 38 | 37 / 38 | | | 37 / 38 | |
| *Streptococcus termophilus* | MRS | 6 / 11 | 4 / 11 | | | 4 / 11 | |
| *Lactobacillus sakei* | MRS | 4 / 7 | 4 / 7 | | | 4 / 7 | |
| *Lactobacillus plantarum* | MRS | 0 / 18 | 0 / 18 | | | 0 / 18 | |
| *Lactobacillus fermentum* | MRS | 0 / 7 | 0 / 7 | | | 0 / 7 | |
| *Streptococcus infantarius* subsp. *infantarius* | MRS | 0 / 2 | 0 / 2 | | | 0 / 2 | |
| *Lactobacillus mucosae* | MRS | 0 / 2 | 0 / 2 | | | 0 / 2 | |
| *Salmonella* spp. | BHI | 0 / 8 | 0 / 8 | | | 0 / 8 | |
| *Staphylococcus aureus* | BHI | 0 / 14 | 0 / 14 | | | 0 / 14 | |
| *Staphylococcus epidermidis* | BHI | 0 / 4 | 0 / 4 | | | 0 / 4 | |
| *Leuconostoc mesenteroides* | MRS | 0 / 9 | 0 / 9 | | 0 / 9 | | |
| *Enterococcus mundtii* | MRS | 0 / 1 | 0 / 1 | | | 0 / 1 | |
| *Lactobacillus scurvatus* | MRS | 0 / 3 | 0 / 3 | | | 0 / 3 | |
| *Pediococcus* spp. | MRS | 0 / 6 | 0 / 6 | | | 0 / 6 | |
| *Lactobacillus delbrueckii* | MRS | 0 / 2 | 0 / /2 | | | 0 / 2 | |

Supplementary Table 6. Antibiotic susceptibility of *Pediococcus acidilactici* ST31BZ, *Pediococcus pentosaceus* ST75BZ and *Pediococcus pentosaceus* ST87BZ.

| Antibiotic | μg/disk | ST31BZ | ST78BZ | ST85BZ |
| --- | --- | --- | --- | --- |
|  |  | Diameter inhibition zone (mm) | | |
| chloramphenicol | 30 | 25 | 22 | 25 |
| neomycin | 10 | 18 | 16 | 16 |
| ampicilin | 10 | 0 | 0 | 0 |
| tobramycin | 10 | 0 | 0 | 0 |
| kanamycin | 30 | 0 | 0 | 0 |
| gentamycin | 10 | 0 | 0 | 0 |
| amikacin | 30 | 0 | 0 | 0 |
| oxacillin | 1 | 0 | 0 | 0 |
| vancomycin | 30 | 0 | 0 | 10 |
| bacitracin | 10 | 17 | 12 | 12 |
| ciprofloxacin | 5 | 0 | 0 | 0 |
| ceforoxime | 30 | 26 | 22 | 23 |
| imipenem | 10 | 28 | 28 | 29 |
| clindamycin | 2 | 25 | 24 | 24 |
| erytromycin | 15 | 23 | 22 | 23 |
| cefepime | 30 | 18 | 16 | 15 |
| penicillin | 10 | 27 | 22 | 22 |
| tetracicline | 30 | 0 | 0 | 0 |
| metronidazole | 50 | 0 | 0 | 0 |
| nalidix acid | 30 | 0 | 0 | 0 |
